# Analytical investigation of non-Fickian receptor-mediated endocytosis

**DOI:** 10.1101/808212

**Authors:** Milad Rismanian, Behzad Damirchi

## Abstract

The most important process in which viruses and bioparticles can enter the animal cell is receptor mediated endocytosis. The main propellant of this phenomenon is the attraction between receptor molecules floating in the lipid bilayer membrane of the host cell and ligand molecules on the target. In this paper, the aforementioned phenomenon is investigated analytically assuming that the diffusion model of the receptor molecules in the lipid bilayer is non-Fickian. Penetration time of the target molecule in the host cell shows that there is a critical limit for the target molecule size. The molecules larger than this critical size will experience an oscillatory motion without passing the host cell membrane. For instance, it is found that the maximum diameter for a typical target molecule with dimensionless relaxation time of 5 for receptor diffusion into a cell membrane is 0.22 of the membrane length.

## Introduction

There are different mechanisms regarding penetration of a molecule into a host cell. Ion transition through cell membrane is usually performed aided by protein channels placed in the membrane. This process can be done selectively or non-selectively [1-3]. Large molecules use larger protein channels to be able to penetrate into their host cell which the most familiar one is water channel [4,5]. Cell penetrating peptides permeate into cell membrane via water pore formation on the membrane [6]. Some of the larger molecules will penetrate via endocytosis process in which the attraction between target molecule ligands and membrane receptors causes the molecule to be surrounded by cell membrane [7] and get absorbed by the cell. Molecular dynamics simulations can reveal detailed observations at this scale. The application of this technique on studies such as assembly of high density proteins (HDL) [7-8], penetration of peptides into living cell [6] and homo-oligomerization of integrin proteins on the membrane [9] have showed the ability of this method to model and further understanding of nano-scale biological processes.

Receptor-mediated endocytosis is the most important mechanism for virus penetration into a cell. After penetration of viruses, they will affect the cell and exit via a similar mechanism named exocytosis. Echavarria-Heras et al. [11] proposed a theoretical model to investigate the effect of mechanisms that influence surface aggregation patterns of LDL receptors near coated pits. Size effect of target molecule in endocytosis is the subject of many researches [12–19]. Prabha et al. [11] examined endocytosis of PLGA nanoparticles through cell membrane experimentally and found that smaller nanoparticles (70 nm) have higher transfection than the larger ones (200 nm) in different cell lines. Osaki et al. [11] suggested the optimal size of approximately 50 nm for cellular uptake. Gao et al. [20] and shi et al. [21] proposed some models for concentration distribution in the endocytosis process. They considered Fick’s law for receptor diffusion in a membrane and solved it for infinite membrane length analytically and for finite length numerically. They suggested that the results of finite length model are closer to real phenomenon. Rismanian et al.[22] showed that Fick’s law cannot predict the diffusion phenomenon in the cell length and time scales. They proposed a lagged model in which there is a phase lag between mass flux and its producing agent i.e. concentration gradient for mass transport in the cell scale for biological phenomena.

In this study receptor-mediated endocytosis of a target molecule into a host cell is investigated analytically. A non-Fickian lagged model and the finite length for the cell membrane are considered. The results showed that molecules larger than a certain size cannot penetrate to the cell. The value of this threshold -A_cr_-is related to the lag time of receptor diffusion into membrane cell. This finding can help pharmacologists to choose optimum size of drugs to penetrate into a definite cell and not penetrate to others.

### Mathematical modeling

Fig.1(a) shows the change of the receptor molecules concentration on the host cell membrane due to descending of the target cells. Before contact with the target molecules, the receptors are assumed to be distributed uniformly on the cell membrane in the state of maximum entropy with constant concentration of *ξ*_0_. After contacting, the receptor concentration increases to the level of ligand concentration *ξ*_L_So, the concentration along cell membrane can be divided into two parts; the first part, 0 < *s < a*(*t*), where the concentration assumed to be *ξ*_L_ and the second part, *a*(*t*) < *s < L*, where the concentration changes along the host membrane length. The a(t) is half of the contacting area of ligand-receptor which is increasing with time from 0 to half of the target molecule circumference and L is half of the host membrane length. The receptors in the vicinity of the contact region are drawn by diffusion, resulting in a local decrease in receptor concentration in that region.

**Fig. 1.**
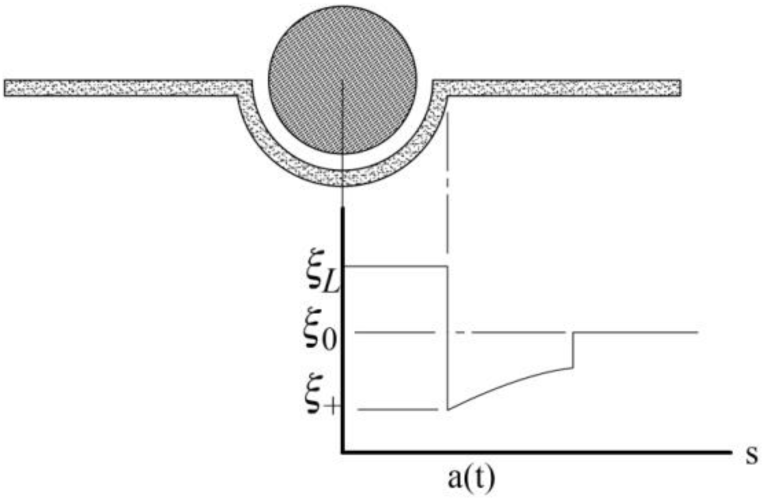
(a) The penetration of the target in the host cells (the Endocytosis phenomenon), (b) The concentration of the receptor molecules along the host cell membrane

The diffusive process of receptors toward the binding site can be characterized by anon-uniform receptor distribution function *ξ* (*s, t*) (Fig.1(b)). The more time passes, the size of the contact area 2a(t) increases.

Considering the continuity and non-Fickian mass transport equations (Eq. (1), (2)), the governing equation of the receptor molecules diffusion in the second part of 2D configuration of the cell membrane can be obtained as Eq. (3).

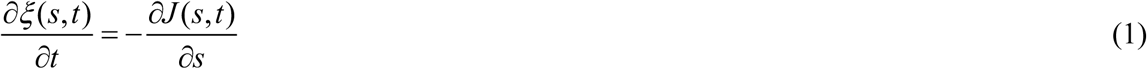

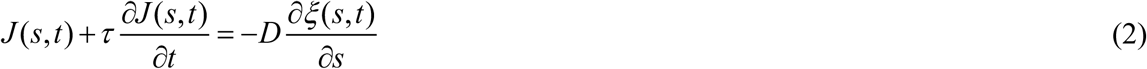

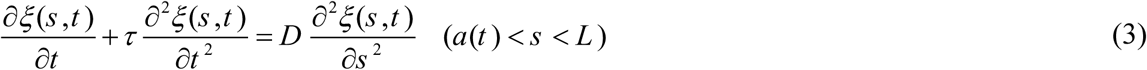

where *J*, *τ*, D represent the mass flux of receptor, the relaxation lag time and receptor diffusion coefficient through the membrane, respectively. The relaxation lag time in non-Fickian mass transport equation is related to the speed of mass transport 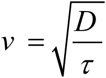.

The boundary and initial conditions for Eq. (3) are assumed through Eqs. (4-7):

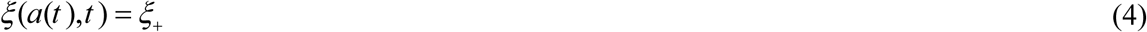

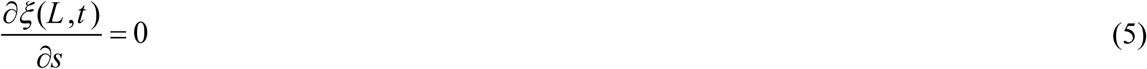

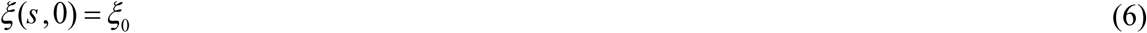

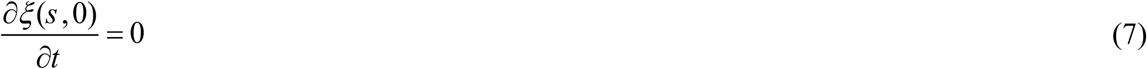

where *ξ*_+_ denotes concentration directly in front of the contact edge.

### Analytical Solution

We consider dimensionless variables as follows.

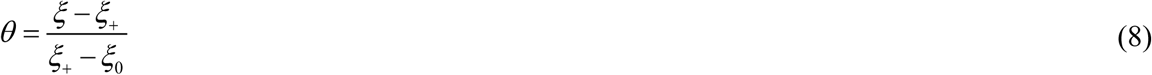

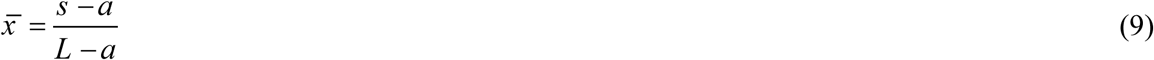

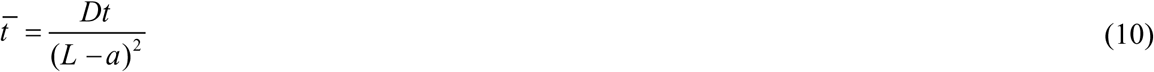

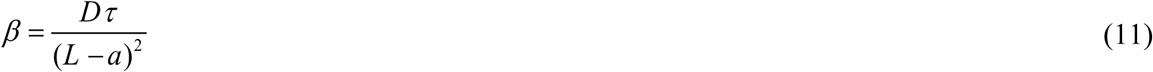

The non-dimensional form of the governing equation and its initial and boundary conditions are:

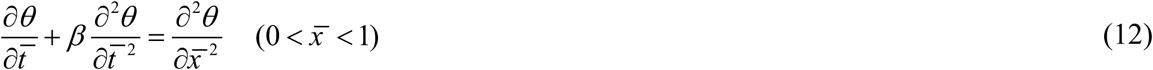

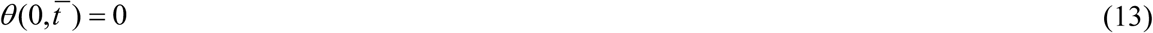

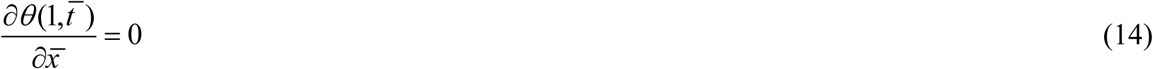

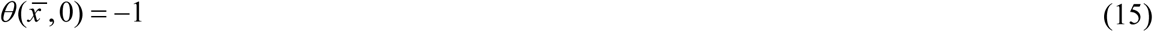

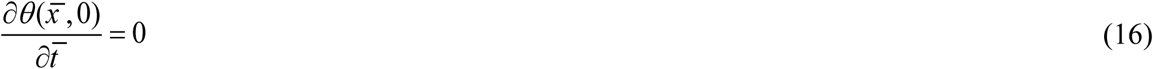

The dimensionless concentration of receptor molecules along the host cell membrane can be obtained with the method of separation of variables as Eq. (17).

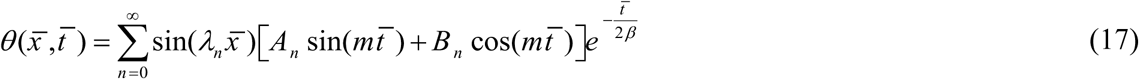

Where corresponding eigen-values are 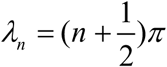 and other variables can be expressed by Eqs. (18-20)

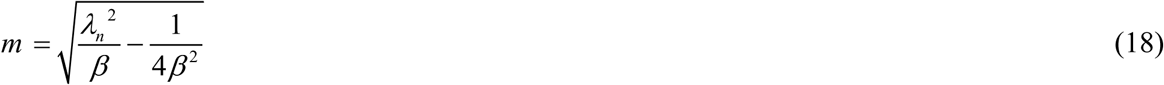

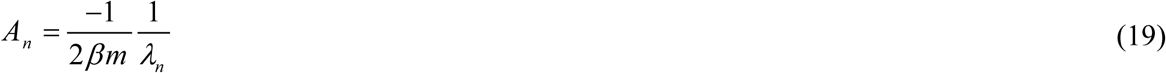

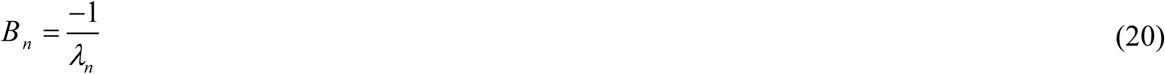

In order to investigate the problem, the two parameters of *a*(*t*) and *ζ* _+_ are desired to be found. These parameters can be obtained using two models as below.

### Equation of Membrane thermal fluctuations

In order to find *ζ*_+_, thermal fluctuations of the membrane should be investigated. Freund and Lin [23] considered the free energy equation for a curved cell membrane in contact with a substrate as Eq. (21).

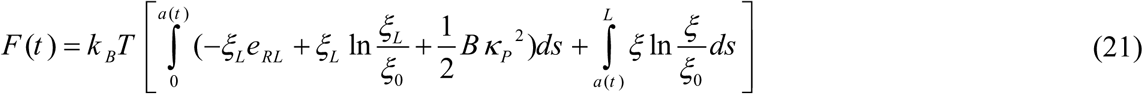

Where k_B_ is the Boltzmann constant, T is the absolute system temperatureis the energy *k*_*B*_*Te*_*RL*,_ of a single receptor–ligand bond, *k*_*B*_*T* ln 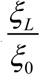 and *k*_*B*_*T* ln 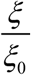 are the entropy loss of bound receptors relative to free ones, respectively [24]. 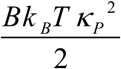 is the elastic bending energy of the membrane wrapping around a cylinder with radius of curvature 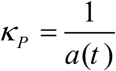 [25]. Thus, the free energy function consists of the energy of receptor–ligand binding, the entropy loss of receptors, and the elastic energy of the cell membrane.

The time derivative of the free energy Eq. (21) is:

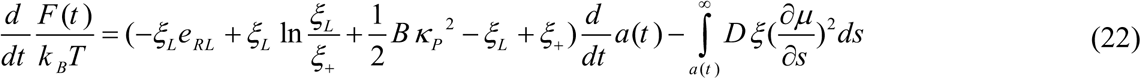

where *μ* is the chemical potential of a receptor defining as bellow,

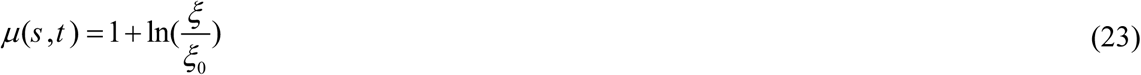

In order to establish a balance between the rate of free energy reduction by wrapping and the rate of energy dissipation by receptor transport [20], the first term on the right hand side of Eq. (22) must be equal to zero so that

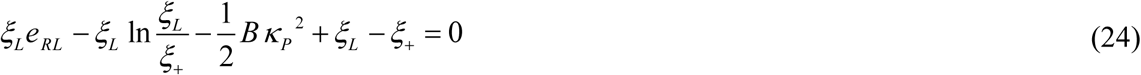

### Conservation of receptor quantity

The total number of receptor on cell membrane is constant during the time. So, a conservation equation can be obtained as Eq. (25)

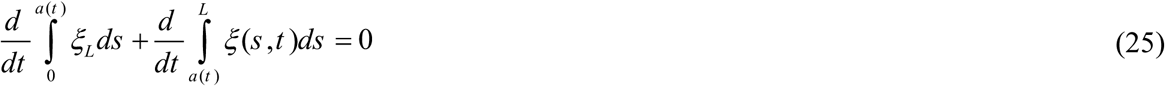

Using the Leibniz rule and Eq. (2), the above equation can be expressed as Eq. (26)

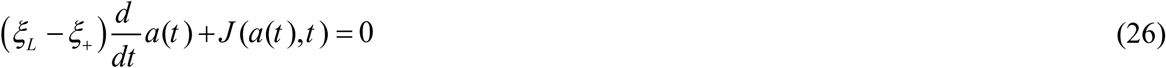

According to Eq. (3), the mass flux of receptor in a(t) is

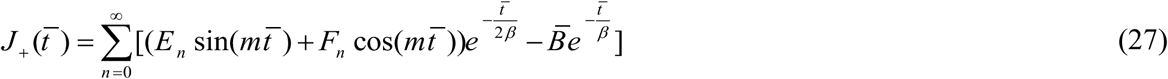

Where:

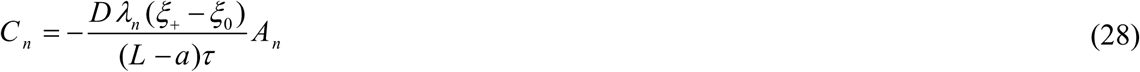

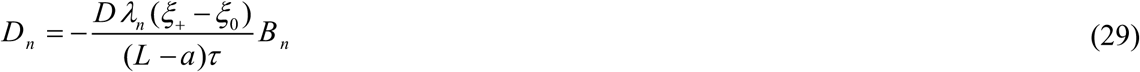

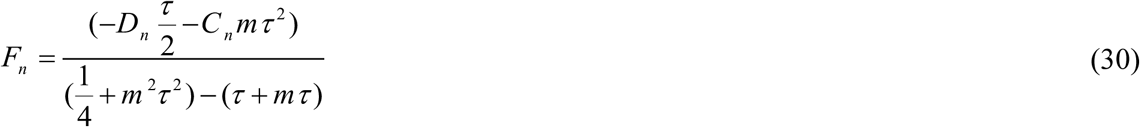

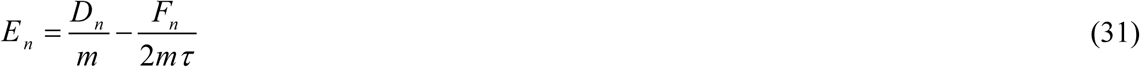

By substituting *ξ*_+_ from Eq. (26) to Eq. (24) we get following relation for rate of a(t) change. Considering initial value of *a*(0) = 0 Eq. (32) can be solved.

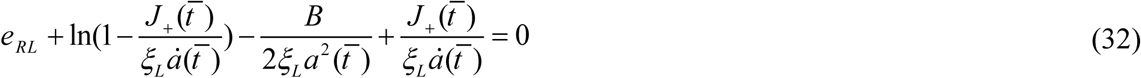

## Results and discussion

The results are investigated for concentration distribution and the penetration velocity for a target molecule with the radius of a. It is expected to have more accurate results rather than available models, because of including finite length of membrane in our analysis. The penetration time is defined by the time in which *a*(*t* _*p*_) = *πR*. The values for *B* / *k*_*B*_*T* and *e*_*RL*_ / *k*_*B*_*T* are set to 20 and 15 similar to other models [4,20,25]. The values of diffusion coefficient, receptor concentration in contact with target molecule and initial concentration of receptors are assumed to be *D =* 1 × 10^–14^ [*m*^2^ / *s*] [26], *ξ*_L_ = 5 × 10^3^ [#/*μm*^2^] and *ξ*_0_ *=* 5 × 10^1^ [#/*μm*^2^] [27–29], respectively. The behavior of the target molecule for different relaxation lag time is investigated to show the effect of lag time on the speed of mass transport.

Fig. 2, shows the profile of dimensionless receptor concentration directly in front of the contact edge as a function of contacting length. As shown, the more contact between target molecule and host cell, the more receptor concentration directly in front of the contact edge. Once the contact starts, the values of contact length and consequently receptor concentration in front of the contact edge are equal to zero. Absence of receptors in start of contact causes the diffusion motion of receptors to this region.

**Fig. 2.**
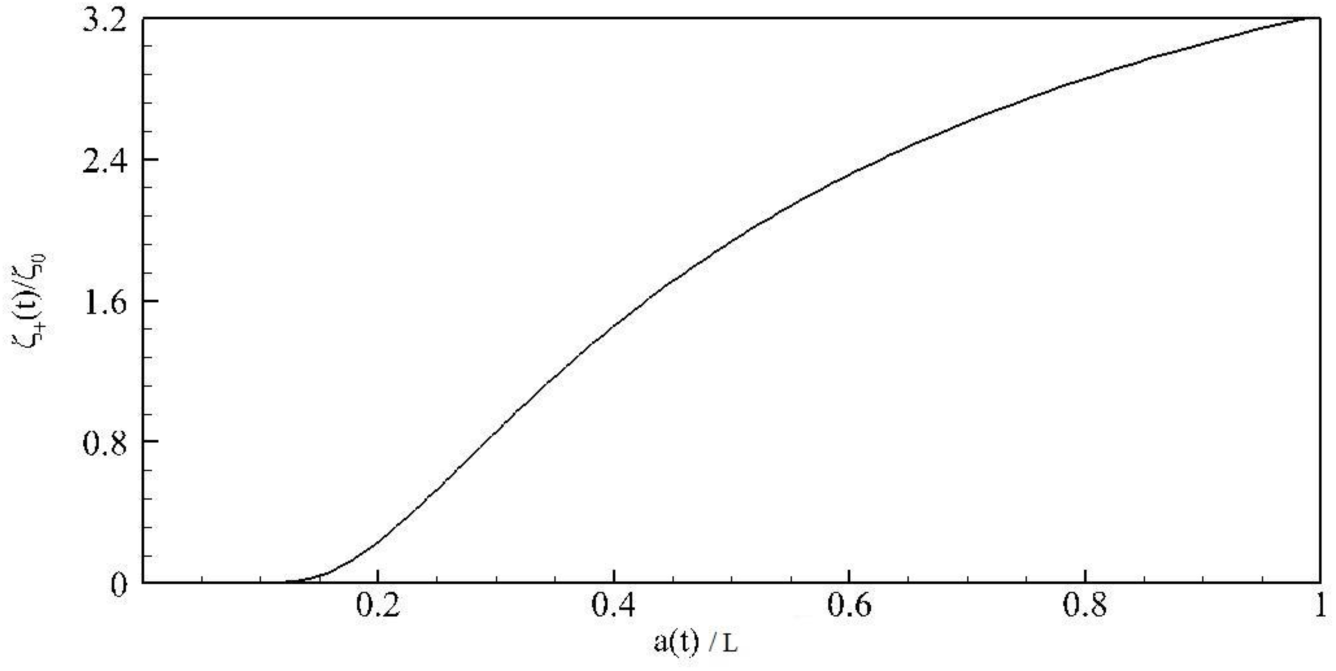
Profile of dimensionless concentration for receptor contacting target molecule directly in front of the contact edge

The profile of dimensionless receptor concentration after contact region is shown in Fig. 3 and Fig. 4 for different non-dimensional times. The difference between these two figures is relaxation time which is selected 5 in Fig. 3 and 0.05 in Fig. 4. Fig. 3a and Fig. 3b predict the place of mass shock in dimensionless time of 2 and 3 according to 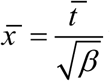 which is in agreement with lag model of mass transport. For 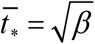, the mass shock reaches to the end of the membrane length and reflected back. After that time, Fig. 3c, the amplitude of the mass shock will decrease. Then after repeated backward and forward motions the shock will be damped.

**Fig. 3.**
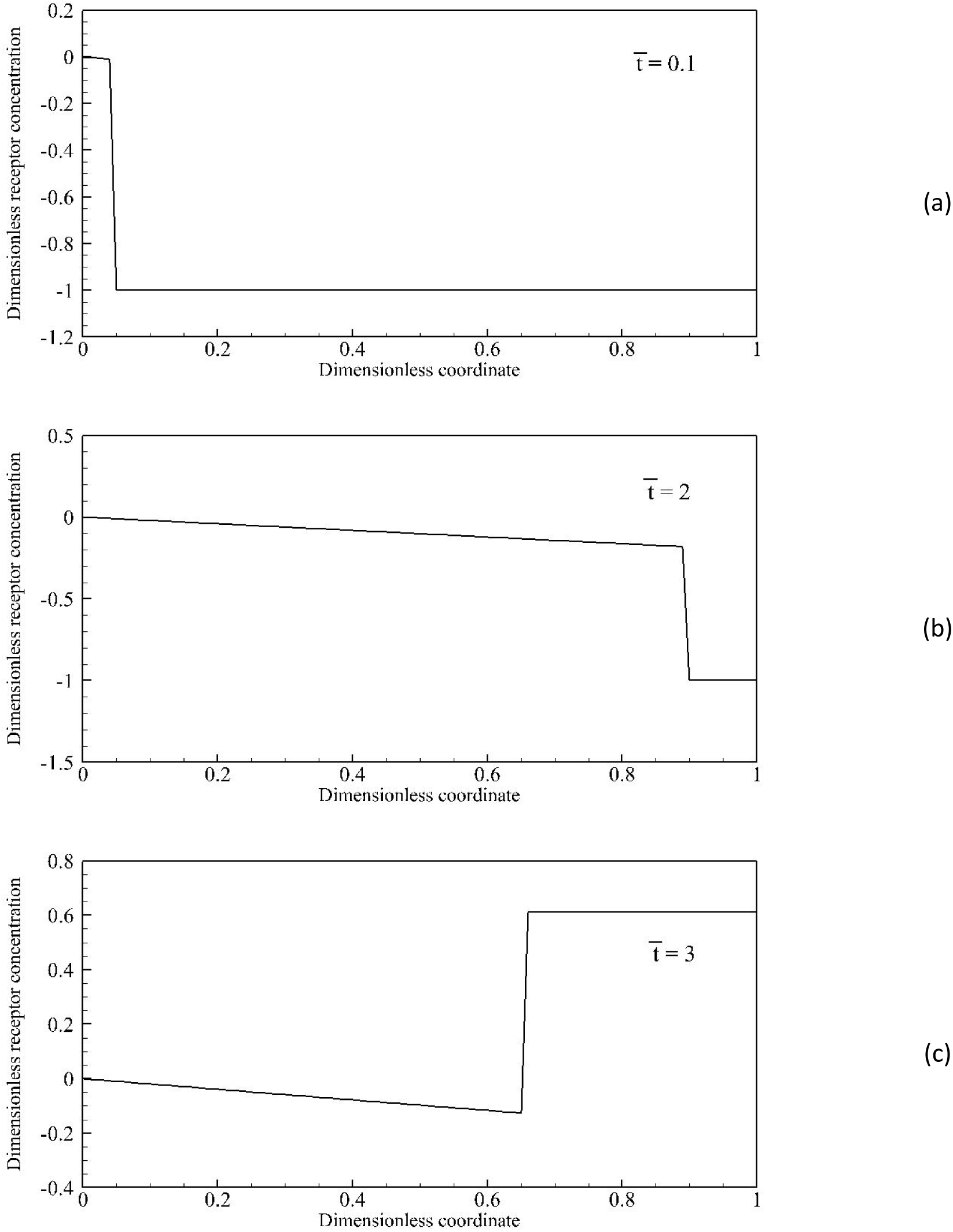
Change of dimensionless concentration for receptor after the contact edge with dimensionless coordinate for dimensionless relaxation time of *β* = 5 ; a) 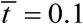; b) 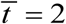; c) 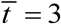

**Fig. 4.**
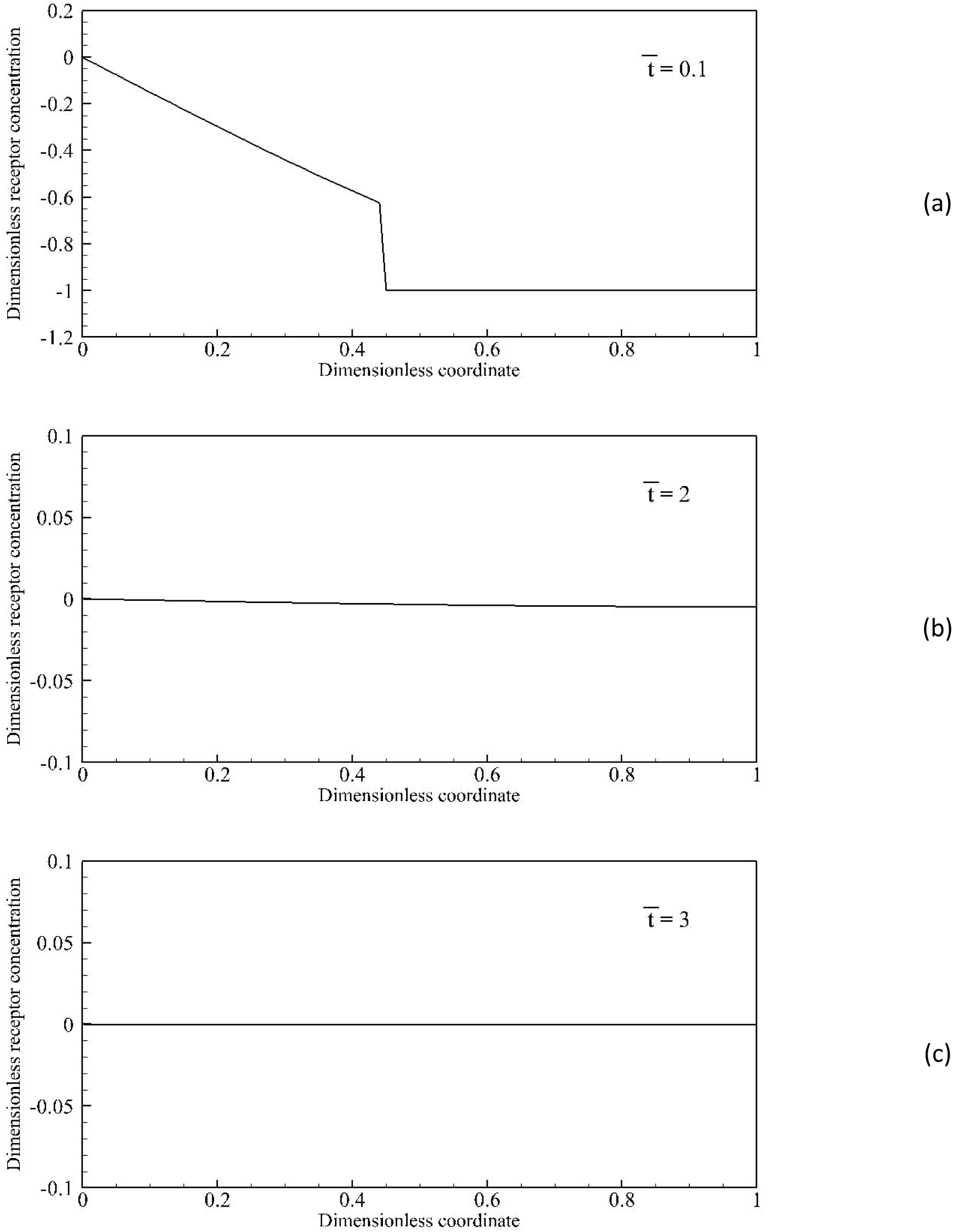
Change of dimensionless concentration for receptor after the contact edge with dimensionless coordinate for dimensionless relaxation time of *β* = 0.05 ; a) 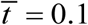; b) 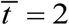; c) 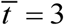

Comparison between Fig. 3 and Fig. 4 shows that in smaller relaxation times, the receptor diffusion behavior obtained from lag model is close to Fick’s model. According to Fig. 4, in small relaxation time for receptors one can use Fick’s model instead of lag model in normal time scales. In contrast, when dealing with phenomena in which time scales are small, using Fick’s model could cause considerable error in predicting actual behavior of the phenomena.

In Fig. 5a, the change of contact length a(t) with penetration time is shown for assumed *β* = 5. According to this figure the value of a(t)/L in this condition cannot exceed 0.34. The penetration time of a molecule is when the a(t) becomes equal to *π D* /2. So, the molecules with D/L larger than 0.22 cannot penetrate to the host cell and experience an oscillatory motion. On the other hand, the smaller molecules will penetrate during certain dimensionless time of 0 to 4.5 as shown in this figure. For example, the penetration time of a molecule with a diameter of 1 micron into a host cell with a length of 10 micron, corresponds to 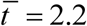 when the a(t)/L reaches 0.16. The results reveal there is an upper limit for the size of biomolecules penetration into host cell, while this phenomena has not seen already. The porpuse of this article is not to obtain an optimum size for biomolecule penetration into host cell. These results show that molecules with radius smaller than the critical size, D_cr_, can penetrate into host cell. It should be noted that in this study *β* is an assumed value and is not according to exact properties of the cell membrane. In order to find the exact value of *β* one should use some extensive molecular dynamics simulations or experimental observations. So, the exact value of this critical size can be obtained considering an exact value for the relaxation time of cell membrane.

**Fig. 5.**
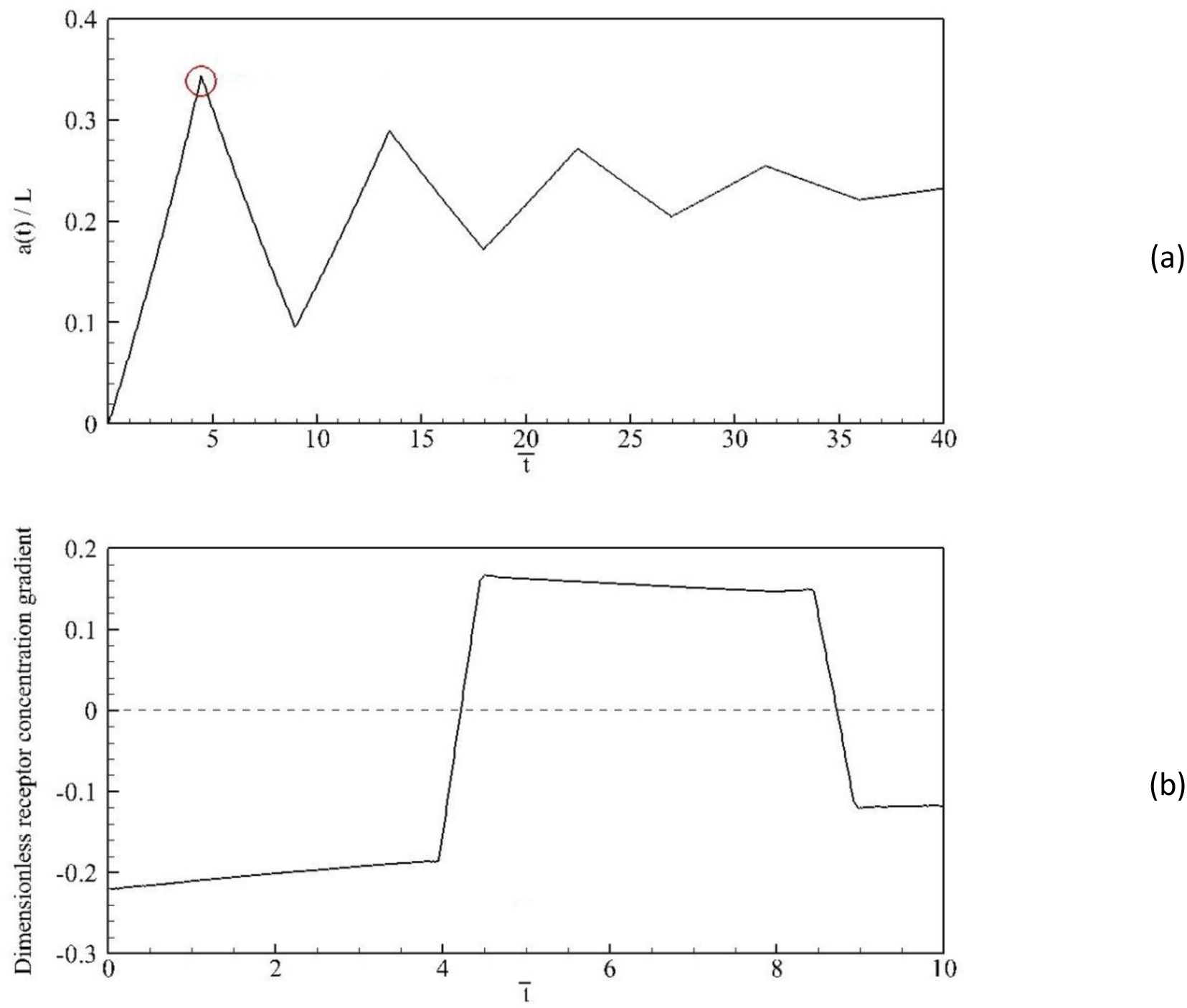
(a) change of contact length a(t) with time; (b) change of gradient of receptor concentration directly in front of the contact length; *β* = 5

It can be deduced that molecules by the surface area larger than A_cr_ will experience an oscillatory motion for entering the cell and finally they could not penetrate the cell. The change of gradient of the receptor concentration directly in front of the contact length is shown in Fig. 5b. Our model predicts that mass flux will oscillatory give positive and negative values and prohibit the larger target molecules from entering the cell via endocytosis. This result is an important finding in drug delivery mechanisms.

## Conclusion

In this paper receptor mediated endocytosis process is investigated assuming that the diffusion model of the receptor molecules in the lipid bilayer is non-Fickian. Different relaxation lag time is chosen to investigate the effect of lag time on the speed of mass transport. Our results show that in smaller relaxation times, the receptor diffusion behavior obtained from lag model is close to Fick’s model. In larger relaxation times, the molecules which their surface are larger than -A_cr_-will oscillate in the cell membrane and could not penetrate the cell. The main objective of our study is to provide an analytical basis helping prediction of the size effect of receptor mediated endocytosis. This prediction may provide some advices for drug delivery systems. The size of the drugs is an important factor in targeting drug delivery for increased efficiency and selectivity.

